# Dysregulated neutrophil extracellular trap formation is a novel driver of clonogenicity and therapeutic vulnerability in CMML

**DOI:** 10.64898/2026.07.28.741351

**Authors:** Saveg Yadav, Callie T. Brown, Mark Cody, William L. Heaton, Claudia de Araujo, Marco Marchetti, Robert A. Campbell, Anthony D. Pomicter, Justin Williams, Christian Con Yost, Shannon E. Elf, Ami B. Patel

## Abstract

Chronic myelomonocytic leukemia (CMML) is an aggressive hematologic malignancy characterized by excess inflammatory signaling and clonal myeloproliferation. The relative contribution of neutrophils (PMNs) to the inflammatory milieu in CMML is poorly understood. In this study we sought to understand whether neutrophil extracellular trap (NET) formation, a key mediator of neutrophilic inflammation, is dysregulated in CMML and can be therapeutically targeted with a novel peptide inhibitor of NETosis called neonatal NET-inhibitory factor (nNIF). Here, we demonstrate that baseline NET formation is aberrantly increased in primary CMML PMNs transcriptionally primed for NETosis, and that soluble factors produced during CMML NET formation promote clonogenicity in CMML CD34^+^ hematopoietic cells matched to the same patient. Further, we show that nNIF and clinical agents under investigation in CMML effectively inhibit NETosis, warranting further study of NET inhibitory agents in this rare disease with limited treatment options.

## Introduction

Neutrophil extracellular traps (NETs) are web-like structures comprised of decondensed chromatin extruded by neutrophil granulocytes (PMNs) as an innate immune response to microbial invasion and tissue damage^1^. The DNA, histones, and antimicrobial proteases released by PMNs during this process of programmed cell death (lytic NETosis) directly promote tissue damage and act as damage-associated molecular patterns (DAMPs) triggering additional inflammatory cascades^2^. Dysregulated NETosis under sterile conditions engenders a cytokine-rich milieu that has been found to drive inflammation, thrombosis and immune perturbations in disease and facilitate cancer cell proliferation and metastasis^3-6^. Chronic myelomonocytic leukemia (CMML) is an aggressive stem cell malignancy distinguished by excess myeloproliferation, hyperactive RAS/MAPK and JAK/STAT signaling, inflammation and autoimmunity^7-9^. While clonal monocytes are well-recognized inflammatory mediators in CMML, the effector functions of clonal PMNs in CMML remain uncharacterized. Considering that NET formation is a major mechanism of neutrophilic inflammation, we sought to interrogate the biology of NETosis in primary CMML cells using neonatal NET inhibitory factor (nNIF), an endogenously occurring peptide previously described by our group that selectively suppresses NET formation without inhibiting critical PMN immunosurveillance functions such as phagocytosis and chemotaxis^10,11^.

## Study design

### Primary samples

Peripheral blood was collected from healthy donors (HD) or CMML patients under University of Utah biobanking protocols (#89989 and #45880). PMNs were isolated using the EasySep Direct Human Neutrophil Isolation Kit (Stemcell Technologies, Cat. #19666) as per the manufacturer’s instructions. Plasma was separated by centrifuging the EDTA-whole blood at 1500xg at 4 °C for 15 minutes.

### NET induction and inhibition

A total of 1 × 10^6^ PMNs were plated on poly-l-lysine (Sigma P1399) coated plates in warmed, serum-free M199 media. PMNs were pretreated with nNIF or its inactive scrambled peptide control (SCR) (10 nM; synthesized at the University of Utah as previously described^10^) for 1 hour and stimulated with either phorbol 12-myristate 13-acetate^1^ (PMA; 10 nM unless otherwise indicated; Sigma, P8139-1MG) or 15% v/v HD or CMML plasma for 2 hours to induce NETs. For drug experiments, HD or CMML PMNs were treated with cobimetinib (0.5-1 µM; Selleck Chem, S804) or ruxolitinib (0.1-0.5 µM; Selleck Chem, INCB18424) alone or in combination for 3 hours. For all experimental conditions, culture supernatants (NET-S) were collected by gentle pipetting. The residual cells and NETs were incubated with 200 µL of micrococcal nuclease (Worthington, LS004797: 185 U/mL) for 15 minutes at room temperature (RT) to release the NETs into solution (NET lysate), and the reaction was quenched with 25 mM EDTA. Samples were stored at -20 °C and analyzed following one freeze-thaw cycle.

### NET quantitation - MPO-DNA complex assay

96-well half area high binding plates (Corning, 3690) were coated with 50 µL of anti-MPO capture antibody (Invitrogen; PA5-16672) per well in coating buffer (Sigma, SRE0034) and incubated overnight at 4 °C. Plates were washed with washing buffer and blocked with 2% bovine serum albumin in PBS for 2 hours. NET lysate (30 µL) or HD/CMML plasma (10 µL) was added. Plates were incubated on a shaker at 300 rpm for 90 minutes. After washing, anti-DNA POD detection antibody (Cell Death Detection ELISA^PLUS^, Sigma 11774425001) was added and incubated on a shaker at 300 rpm for 90 minutes in the dark. After washing, 100 µL of peroxidase substrate solution was added and incubated on a shaker at 300 rpm for 15 minutes in the dark. Reactions were terminated with 50 µL of hydrochloric acid, and absorbance was measured at 450 nm using an Agilent BioTek Synergy H1 microplate reader.

### NET visualization - Immunofluorescence

A total of 2 × 10^5^ PMNs were plated on poly-l-lysine-coated chamber slides. Cells were stained with SYTO Green (1 µM; cell permeable) and SYTOX Orange (1 µM; cell impermeable) for 5 minutes and visualized using an Olympus FV-3000 confocal microscope (10-12 1-μm steps per z-stack image).

### Bulk RNA sequencing and analysis

Peripheral blood PMNs were isolated from HD (n = 5) and CMML patients (n = 7), lysed, and prepared for RNA sequencing. TrimGalore (v0.6.10) was used to remove low quality reads and adapters from raw RNA-seq reads. Filtered reads were aligned to the human genome (GRCh38.113) using STAR aligner (v2.7.11a) with the “--quantMode GeneCounts” option. To assess sample quality, raw gene counts were converted to Counts Per Million, log2-transformed, then used for Principal Component Analysis using the Python package scikit-learn (v1.5.1). The first two principal components were used for plotting samples using the Python packages matplotlib (v3.9.2) and seaborn (v0.13.2). Raw gene counts from STAR were then used for pairwise differential expression analysis using DESeq2 (v1.42.1). Genes with a Benjamini-Hochberg adjusted p-value < 0.05 were considered significantly differentially expressed. A volcano plot was generated with the R package (EnhancedVolcano) for an overview of the differential expression analysis. The top differentially expressed genes were visualized using a heatmap generated with the R package pheatmap (v1.0.13) to compare expression patterns between healthy donor PMNs and CMML PMNs. Differentially expressed genes were then used for pre-ranked Gene Set Enrichment Analysis (GSEA) and over-representation analysis (ORA) of Gene Ontology Biological Processes or Hallmark gene sets. The R package clusterProfiler (v4.16.0) was used for GSEA, while ORA was performed using custom code available on Github (https://github.com/mmarchetti90/project_setup_assistant/blob/main/code_base/enrichment_an_alysis_1.py).

### Colony Forming Unit (CFU) assay

CD34^+^ cells were plated in duplicate at a concentration of 1000 cells/dish in 35 mm culture dishes in methylcellulose-based media (MethoCult™ H4535, StemCell), supplemented with 15% v/v NET-S or plasma. Colonies were manually counted after 14 days with a microscope (Laxco SLI3P-FLP) and images were captured at 10X magnification (SeBa software).

### Statistical Analysis

Statistical analysis was performed using GraphPad Prism v10 software. Statistical significance between two groups was determined using a two-tailed unpaired Student’s t-test. For multiple group comparisons, one-way or two-way analysis of variance (ANOVA) tests were performed, followed by Šídák or Tukey’s post-hoc testing. A *p*-value <0.05 was considered statistically significant. Error bars are depicted as mean ± standard deviation.

## Results and Discussion

### CMML PMNs display aberrant NETosis and NET induction under basal conditions

Systemic inflammation driven by clonal myeloproliferation is a well-established feature of CMML yet the relative contribution of PMNs to the inflammatory state remains ill-defined. We hypothesized that similar to other malignancies, altered NETosis may contribute to the inflammatory milieu of CMML. To test for differences in NET-forming capacity between participants with CMML and healthy donor (HD) controls, baseline MPO-DNA complex levels, a surrogate for NET formation^11,12^, were quantified in vitro in NET lysates following stimulation with PMA (**Fig. 1, A-C**). A significant increase in MPO-DNA complex levels at baseline was noted in CMML patients versus HD. Upon stimulation with PMA, MPO-DNA complex levels were similar between CMML and controls, suggesting that CMML patients are close to maximal NET induction capacity under basal, unstimulated conditions. Increased spontaneous NET formation in CMML PMNs was confirmed qualitatively by live cell confocal microscopy using SYTO Green and SYTOX Orange stains (**Fig. 1D**). Given the enhanced NET-forming capacity observed in CMML PMNs, we tested primary CMML and HD plasma for circulating MPO-DNA complexes (**Fig.1E**) and found levels to be qualitatively increased in CMML, providing evidence of augmented NETosis in vivo. Taken together, these data suggest that NET formation is dysregulated in CMML.

**Fig. 1.**
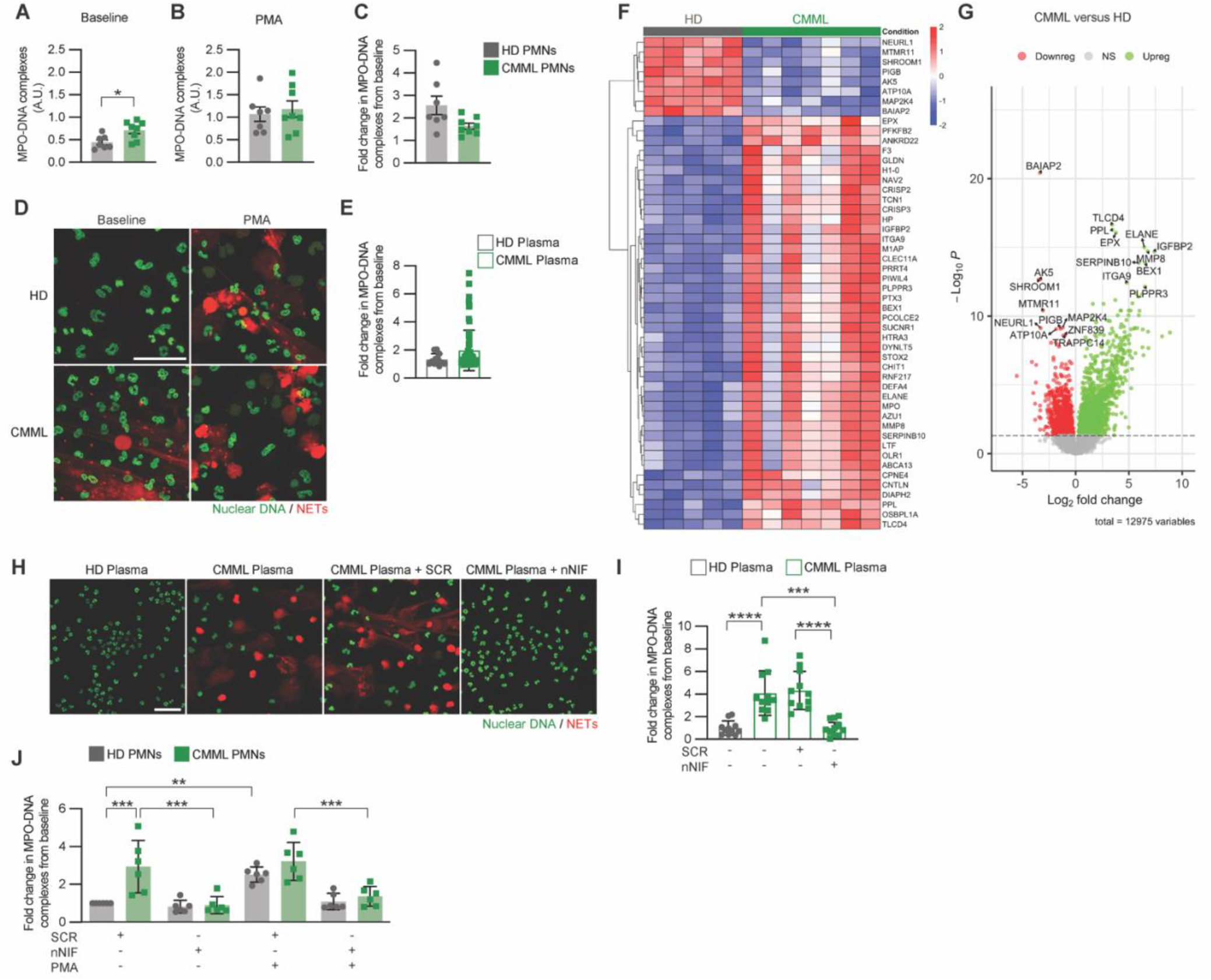
Dysregulated NETosis is a feature of CMML that can be therapeutically targeted with nNIF. (**A**) Quantitation of MPO-DNA complex levels, a surrogate for NET formation, in HD PMNs and CMML at baseline, and (**B**) after stimulation with 50 nM PMA. (**C**) Post-stimulation fold-change in MPO-DNA complex levels. (**D**) Confocal microscopy of HD and CMML PMNs stained with SYTO Green (intracellular DNA) and SYTOX Orange (extracellular DNA). (**E**) Quantification of MPO-DNA complex levels in HD and CMML plasma. (**F**) Heat map and (**G**) volcano plot of top differentially expressed genes in HD versus CMML PMNs (**H-I**) Confocal microscopy and MPO-DNA complex level quantitation in HD PMNs preincubated with nNIF (10 nM) versus SCR (10 nM) peptides followed by incubation with autologous HD or CMML plasma). (**J**) MPO-DNA complex level quantitation in HD and CMML PMNs preincubated with SCR (10 nM) versus nNIF (10 nM) peptides at baseline and following PMA stimulation (10 nM). Data are shown as mean ± SD. Statistical significance was determined by unpaired *t*-tests (**A-C, E**), one-way ANOVA (**I**) or two-way ANOVA (**J**). \**p*<0.05, **<0.01, ***<0.001, ****<0.0001. A.U., arbitrary units (absorbance at 450nm). Scale bar on confocal images = 10µm.

### CMML PMNs exhibit an inflammatory transcriptome that is primed for NETosis

We next asked whether CMML PMNs exhibited baseline transcriptomic changes that could account for their differential NET-forming capacity. To characterize the gene expression profile of CMML PMNs, we performed bulk RNA sequencing on PMNs isolated from HD and CMML participants. The top differentially expressed genes (Benjamini-Hochberg adjusted p-value < 0.05) are depicted in **Fig. 1F-G**. Significantly upregulated genes in CMML PMNs included multiple cytotoxic proteins involved in cell division, primary and secondary granule biosynthesis, inflammation and NETosis, including neutrophil elastase (*ELANE*), myeloperoxidase (*MPO*), tissue factor (*F3*), pentraxin 3 protein (*PTX3*), chitinase 1 (*CHIT1*), neutrophil defensin 4 (*DEFA4*), azurocidin (*AZU1*), neutrophil collagenase (*MMP8*) and lactoferrin (*LTF*). Expression of these genes is known to occur in early granulopoiesis, as myeloblasts differentiate to promyelocytes and myelocytes while retaining mitotic capability^13^. Gene set enrichment analysis demonstrated significant enrichment in pathways regulating cell division and proliferation, neutrophil activation, chemotaxis, oxidative and pathogen-induced stress, and TNF-α signaling via NF-κB in CMML PMNs. These data support that CMML PMNs are activated and exhibit an immature granulocytic transcriptional program characteristic of inflammatory states such as sepsis that trigger emergency granulopoiesis^14-18^, and, in conjunction with the enhanced NET induction observed in CMML PMNs in the absence of agonist therapy, support that CMML PMNs are exquisitely primed for NET formation.

### CMML plasma induces NET formation in healthy PMNs that is blocked by nNIF

Our previous finding of increased MPO-DNA complexes in CMML versus HD plasma led us to ask whether CMML plasma also contains factors that stimulate NETosis in healthy PMNs. To answer this, MPO-DNA complex levels were quantified in NET lysates from HD PMNs incubated with autologous HD or CMML patient plasma (**Fig. 1H-I**). MPO-DNA complex levels significantly increased in HD PMNs following incubation with CMML plasma compared to HD plasma. HD MPO-DNA complex levels following incubation with CMML plasma were completely suppressed by preincubation of HD PMNs with nNIF, an endogenous NET-inhibitory peptide, but unaffected by SCR, its inactive control. These results demonstrate that exposure to CMML plasma stimulates NETosis in healthy PMNs that is prevented by nNIF exposure.

### nNIF prevents basal NETosis in CMML PMNs and inhibits NET induction by PMA in healthy PMNs

To investigate whether nNIF inhibits NET formation in CMML PMNs, primary HD and CMML PMNs were isolated and preincubated with nNIF or SCR. nNIF treatment significantly decreased MPO-DNA complex levels in NET lysates from CMML PMNs both in the presence and absence of PMA (**Fig. 1J**). In addition, nNIF significantly decreased MPO-DNA complex levels in NET lysates from HD PMNs stimulated with PMA, while HD MPO-DNA complex levels in the absence of PMA were comparable between SCR and nNIF treatment groups, as expected. These data show that nNIF treatment effectively prevents NET formation in CMML PMNs.

### Healthy and CMML CD34^+^ cell proliferation is enhanced by factors within CMML NET-S, with nNIF treatment preventing these effects

Aberrant NETosis has been linked to cancer cell proliferation in many tumors. We hypothesized that inflammatory mediators released during NET formation in CMML may have direct effects on hematopoietic stem and progenitor cell (HSPC) growth. Given our previous finding that CMML plasma contains MPO-DNA complexes, we first tested the effect of adding CMML or HD plasma (15% v/v) to methylcellulose-based media in colony-forming unit (CFU) assays using HD CD34^+^ HSPCs. We found that healthy CD34^+^ HSPC growth was significantly enhanced by addition of CMML plasma compared to HD plasma (**Fig. 2A-B**), suggesting that byproducts of NETosis contained within CMML plasma may promote HSPC proliferation. To more directly link dysregulated CMML NET activity to HSPC proliferation, CFU assays were performed with HD CD34^+^ HSPCs in the presence or absence of 15% v/v NET assay supernatant (NET-S; added to methylcellulose-based media) collected from autologous HD or CMML PMNs cultured with SCR or nNIF in serum-free media (**Fig. 2C-D**). Compared to HD CD34^+^ colonies supplemented with HD NET-S, HD CD34^+^ colonies supplemented with CMML NET-S exhibited larger colony size and significantly increased colony number. Treatment of CMML PMNs with nNIF reduced CMML NET-S-treated HD CD34^+^ colony numbers to those observed with HD NET-S **(2D)**. These results demonstrate that soluble factors produced during NETosis in CMML directly contribute to HD CD34^+^ HSPC proliferation and that this effect on CD34^+^ cell growth can be inhibited by blocking NETosis in CMML PMNs using nNIF.

**Fig. 2.**
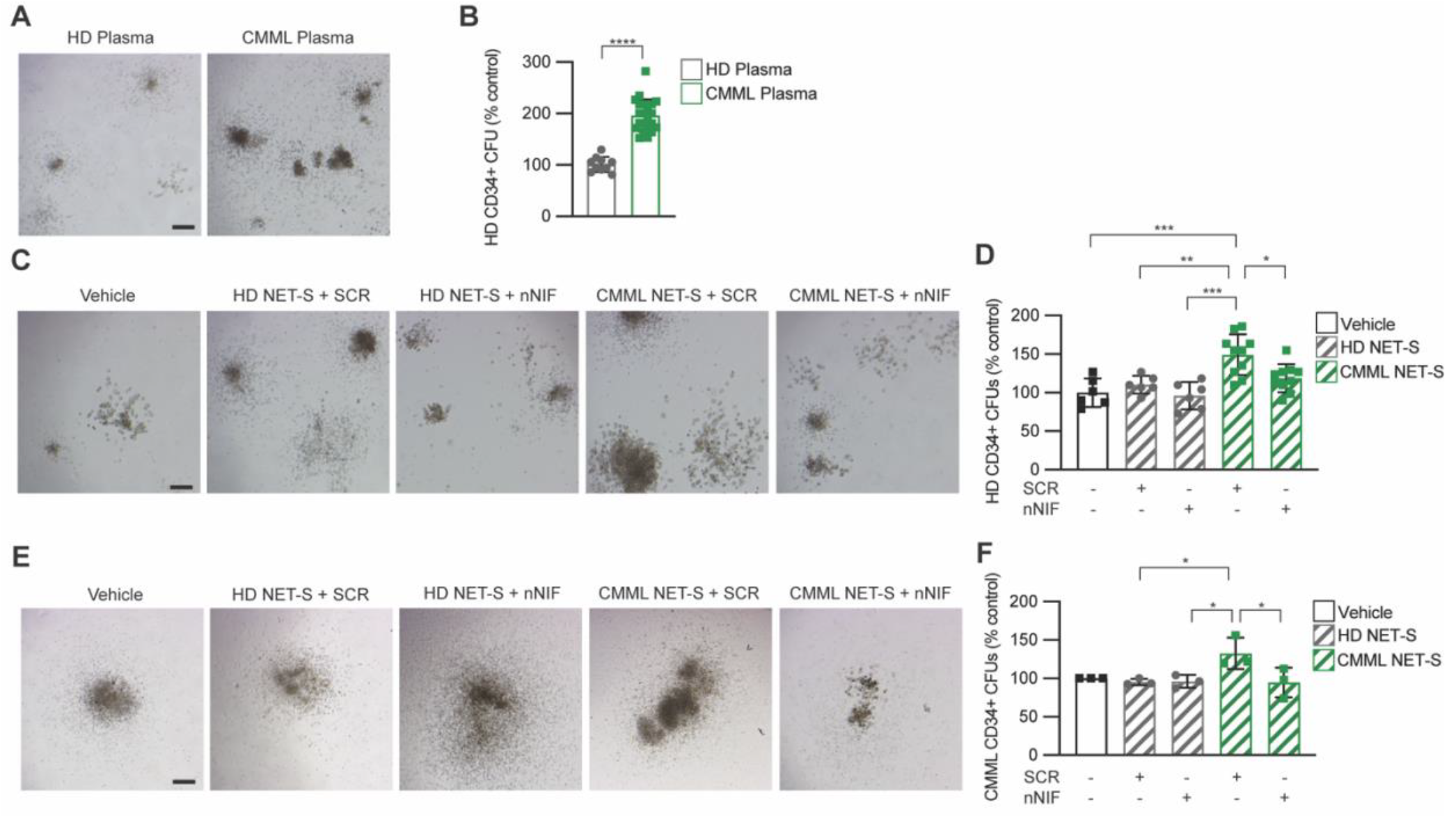
CMML NET-S promotes clonogenicity of HD and CMML CD34^+^ cells that is inhibited by nNIF treatment of PMNs. (**A, B**) Size and quantitation of HD CD34^+^ CFUs cultured with 15% HD or CMML plasma. (**C, D**) Size and quantitation of HD CD34^+^ CFUs cultured with 15% NET-S from HD or CMML PMNs preincubated with SCR (10 nM) versus nNIF (10 nM) peptides. (**E, F**) CMML CD34^+^ CFUs cultured with 15% v/v NET-S from HD or CMML PMNs preincubated with SCR (10 nM) versus nNIF (10 nM) peptides. Data are shown as mean ± SD. Statistical significance was determined by unpaired *t*-tests (**B**), or one-way ANOVA (**D, E**). \**p*<0.05,**<0.01, ***<0.001, ****<0.0001. Scale bar = 10 µm.

To determine whether CMML NET-S had direct proliferative effects on CD34^+^ leukemia stem and progenitor cell (LSPC) proliferation, identical colony assays were performed using primary CD34^+^ CMML cells. CMML CD34^+^ cells were cultured in methylcellulose-based media in the presence or absence of 15% v/v NET-S from HD or autologous CMML PMNs treated with nNIF or SCR. NET-S from autologous CMML PMNs treated with SCR significantly increased CMML CD34^+^ colony number over NET-S from HD PMNs treated with SCR (**2E-F**). NET-S from nNIF-treated CMML PMNs blocked the increase in CD34^+^ CMML colony number observed with NET-S from SCR-treated CMML PMNs **(2F)**. These data show that like HD CD34^+^ cells, CMML CD34^+^ cell proliferation is augmented by autologous CMML NET-S and that LSPC growth can be suppressed by preventing NETosis of CMML PMNs using nNIF. Together, these data provide compelling evidence that CMML NET formation enhances clonogenicity of CMML LPSCs and suggest that targeting NETosis in CMML may exert disease-modifying effects at the stem cell level.

### JAK1/2 and MEK inhibition reduce NETosis in CMML

Agents under active investigation in CMML include the approved inhibitors ruxolitinib (NCT03722407) and cobimetinib (NCT04409639). JAK1/2 inhibitors are recognized for their anti-inflammatory effects and have been previously described to inhibit NETosis and thrombosis in myeloproliferative neoplasms^3^. More recently, MEK inhibitors have been reported to suppress inflammatory responses, and RAS/MAPK pathway activation is a key inducer of NET formation^19,20^. We tested these clinical inhibitors alone and in combination to determine whether they could effectively prevent spontaneous NET formation in CMML PMNs (**Fig. 3**). While both inhibitors significantly reduced CMML PMN MPO-DNA complex levels in NET lysates as single agents, only the combination reduced CMML MPO-DNA-complex levels to that observed with HD PMNs.

**Fig. 3.**
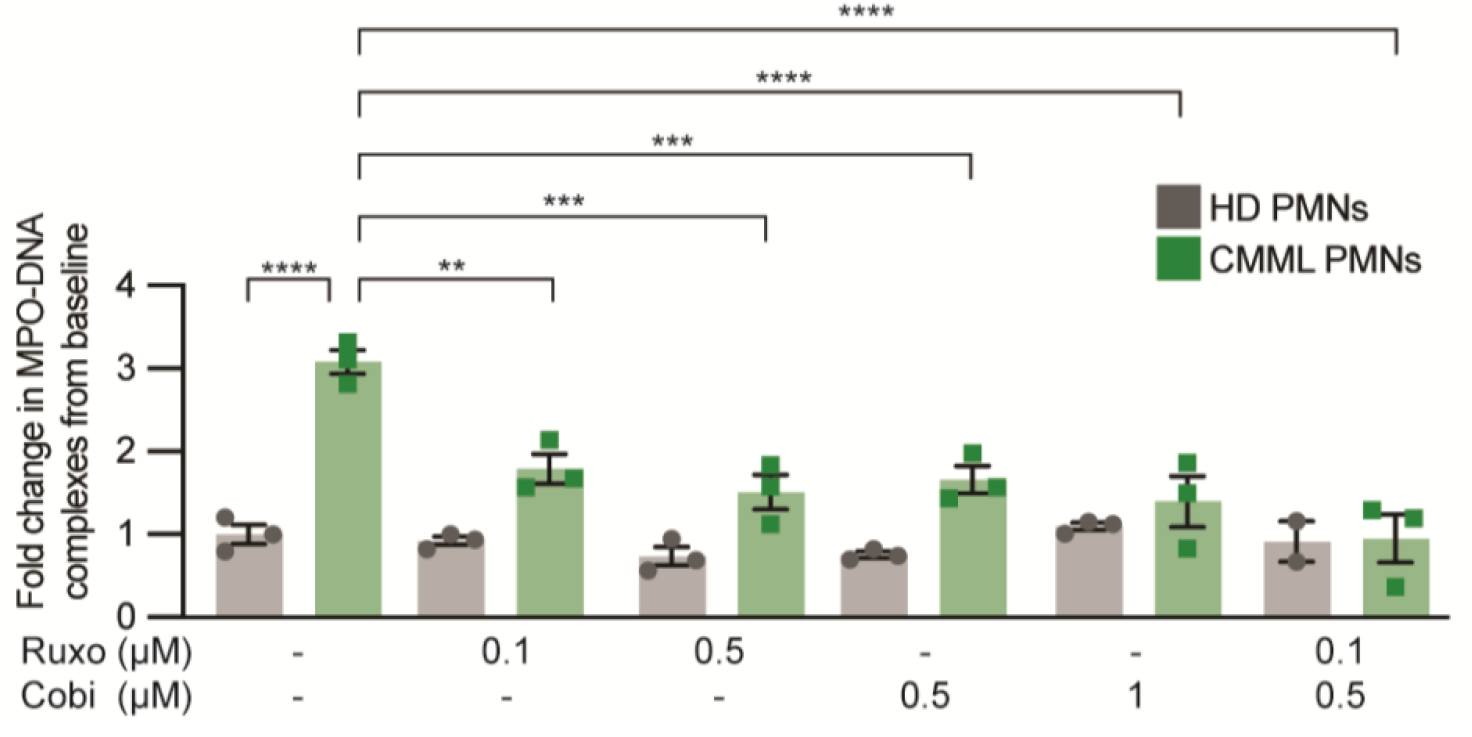
Combined JAK1/2 and MEK inhibition most effectively reduces MPO-DNA complex levels, a surrogate for NET formation, in CMML. MPO-DNA complex level quantitation in HD or CMML PMNs incubated with ruxolitinib (0.1 µM or 0.5 µM) and cobimetinib (0.5 µM or 1 µM), alone and in combination. Data are shown as mean ± SD. Statistical significance was determined by two-way ANOVA. *p < 0.05, **p < 0.01, ***p < 0.001, ****p < 0.0001.

Our results demonstrate that clonal NETosis is aberrantly increased in CMML and that CMML PMNs are maximally primed for NET formation at baseline. CMML plasma induces NET production in healthy PMNs, suggesting that non-neoplastic PMNs in CMML may also contribute to systemic inflammation via NETosis. We show that soluble factors contained within CMML NET-S directly promote both HD and CMML CD34^+^ cell proliferation. These data suggest that neutrophilic inflammation in CMML exerts selective pressure on both non-neoplastic HSPCs and CMML CD34^+^ LSPCs and may contribute to CMML disease maintenance and clonal evolution driving disease progression. Importantly, this work demonstrates that CMML PMN treatment with nNIF, a highly specific inhibitor of NETosis, effectively prevents both baseline clonal NETosis and stimulated HD NETosis, and protects against the proliferative effects of CMML NET-S on healthy and CMML CD34^+^ cell growth and expansion, suggesting that inhibition of NETosis may reduce clonogenicity in CMML. Finally, we demonstrate that clinical agents under investigation for CMML, including ruxolitinib and cobimetinib, recapitulate the effect of nNIF treatment in CMML PMNs, intimating that a component of their anti-inflammatory and anti-proliferative efficacy in CMML is mediated through NET inhibitory activity and suppression of neutrophilic inflammation.

## Acknowledgements

The authors acknowledge the research contributions of participants with CMML and healthy volunteers at the Huntsman Cancer Institute to this study.

A.B.P. was supported in part by the American Society of Hematology (Scholar Award), the Huntsman Cancer Institute (5 For the Fight fellowship), and the NCI (K08CA252883).

S.E.E. was supported by the NIH (C.C.Y. was supported by the NICHD (R01HD093826).

## Authorship contributions

Study conceptualization: A.B.P.

Study methodology, data generation and analysis: S.Y., C.T.B., M.C., W.H., C.A., M.M., R.C., C.C.Y., S.E.E., A.B.P

Patient sample acquisition, compliance, inventory curation: A.D.P, J.A.W. Funding: A.B.P., S.E.E., C.C.Y.

Manuscript writing, reviewing, editing: all authors

## References

1. Brinkmann, V., Reichard, U., Goosmann, C., Fauler, B., Uhlemann, Y., Weiss, D.S., Weinrauch, Y., and Zychlinsky, A. (2004). Neutrophil extracellular traps kill bacteria. Science 303, 1532–1535. 10.1126/science.1092385.

2. Papayannopoulos, V. (2018). Neutrophil extracellular traps in immunity and disease. Nat Rev Immunol 18, 134–147. 10.1038/nri.2017.105.

3. Wolach, O., Sellar, R.S., Martinod, K., Cherpokova, D., McConkey, M., Chappell, R.J., Silver, A.J., Adams, D., Castellano, C.A., Schneider, R.K., et al. (2018). Increased neutrophil extracellular trap formation promotes thrombosis in myeloproliferative neoplasms. Sci Transl Med 10. 10.1126/scitranslmed.aan8292.

4. Yazdani, H.O., Roy, E., Comerci, A.J., van der Windt, D.J., Zhang, H., Huang, H., Loughran, P., Shiva, S., Geller, D.A., Bartlett, D.L., et al. (2019). Neutrophil Extracellular Traps Drive Mitochondrial Homeostasis in Tumors to Augment Growth. Cancer Res 79, 5626–5639. 10.1158/0008-5472.CAN-19-0800.

5. Nie, M., Yang, L., Bi, X., Wang, Y., Sun, P., Yang, H., Liu, P., Li, Z., Xia, Y., and Jiang, W. (2019). Neutrophil Extracellular Traps Induced by IL8 Promote Diffuse Large B-cell Lymphoma Progression via the TLR9 Signaling. Clin Cancer Res 25, 1867–1879. 10.1158/1078-0432.CCR-18-1226.

6. Adrover, J.M., McDowell, S.A.C., He, X.Y., Quail, D.F., and Egeblad, M. (2023). NETworking with cancer: The bidirectional interplay between cancer and neutrophil extracellular traps. Cancer Cell 41, 505–526. 10.1016/j.ccell.2023.02.001.

7. Marando, L., Csizmar, C.M., and Patnaik, M.M. (2025). Chronic myelomonocytic leukemia: molecular pathogenesis and therapeutic innovations. Haematologica 110, 22–36. 10.3324/haematol.2024.286061.

8. Patel, A.B., and Deininger, M.W. (2021). Genetic complexity of chronic myelomonocytic leukemia. Leuk Lymphoma 62, 1031–1045. 10.1080/10428194.2020.1856837.

9. Patel, A.B., Pettijohn, E.M., Abedin, S.M., Raps, E., and Deininger, M.W. (2019). Leukemoid reaction in chronic myelomonocytic leukemia patients undergoing surgery: perioperative management recommendations. Blood Adv 3, 952–955. 10.1182/bloodadvances.2019032300.

10. Yost, C.C., Schwertz, H., Cody, M.J., Wallace, J.A., Campbell, R.A., Vieira-de-Abreu, A., Araujo, C.V., Schubert, S., Harris, E.S., Rowley, J.W., et al. (2016). Neonatal NET-inhibitory factor and related peptides inhibit neutrophil extracellular trap formation. J Clin Invest 126, 3783–3798. 10.1172/JCI83873.

11. Campbell, R.A., Campbell, H.D., Bircher, J.S., de Araujo, C.V., Denorme, F., Crandell, J.L., Rustad, J.L., Monts, J., Cody, M.J., Kosaka, Y., and Yost, C.C. (2021). Placental HTRA1 cleaves alpha1-antitrypsin to generate a NET-inhibitory peptide. Blood 138, 977–988. 10.1182/blood.2020009021.

12. Aoki, J.A., Denorme, F., Cody, M.J., Perry, D.P., Rustad, J.L., Brown, S.M., Goldstein, S.A., Middleton, E.A., Yost, C.C., Harris, E.S., et al. (2025). Plasma surrogate markers of neutrophil extracellular traps correlate with disease severity in patients with moderate to severe acute respiratory distress syndrome. J Inflamm (Lond) 22, 22. 10.1186/s12950-025-00448-8.

13. Lawrence, S.M., Corriden, R., and Nizet, V. (2018). The Ontogeny of a Neutrophil: Mechanisms of Granulopoiesis and Homeostasis. Microbiol Mol Biol Rev 82. 10.1128/MMBR.00057-17.

14. Velasquez, S.Y., Coulibaly, A., Sticht, C., Schulte, J., Hahn, B., Sturm, T., Schefzik, R., Thiel, M., and Lindner, H.A. (2022). Key Signature Genes of Early Terminal Granulocytic Differentiation Distinguish Sepsis From Systemic Inflammatory Response Syndrome on Intensive Care Unit Admission. Front Immunol 13, 864835. 10.3389/fimmu.2022.864835.

15. Calzetti, F., Finotti, G., Tamassia, N., Bianchetto-Aguilera, F., Castellucci, M., Cane, S., Lonardi, S., Cavallini, C., Matte, A., Gasperini, S., et al. (2022). CD66b(-)CD64(dim)CD115(-) cells in the human bone marrow represent neutrophil-committed progenitors. Nat Immunol 23, 679–691. 10.1038/s41590-022-01189-z.

16. Montaldo, E., Lusito, E., Bianchessi, V., Caronni, N., Scala, S., Basso-Ricci, L., Cantaffa, C., Masserdotti, A., Barilaro, M., Barresi, S., et al. (2022). Cellular and transcriptional dynamics of human neutrophils at steady state and upon stress. Nat Immunol 23, 1470–1483. 10.1038/s41590-022-01311-1.

17. Guenther, T., Coulibaly, A., Velasquez, S.Y., Schulte, J., Fuderer, T., Sturm, T., Hahn, B., Thiel, M., and Lindner, H.A. (2024). Transcriptional pathways of terminal differentiation in high- and low-density blood granulocytes in sepsis. J Inflamm (Lond) 21, 40. 10.1186/s12950-024-00414-w.

18. Deschamps, P., Wacheux, M., Gosseye, A., Morabito, M., Pages, A., Lyne, A.M., Alfaro, A., Rameau, P., Imanci, A., Chelbi, R., et al. (2024). CXCL8 secreted by immature granulocytes inhibits WT hematopoiesis in chronic myelomonocytic leukemia. J Clin Invest 134. 10.1172/JCI180738.

19. Stieglitz, E., Lee, A.G., Angus, S.P., Davis, C., Barkauskas, D.A., Hall, D., Kogan, S.C., Meyer, J., Rhodes, S.D., Tasian, S.K., et al. (2024). Efficacy of the Allosteric MEK Inhibitor Trametinib in Relapsed and Refractory Juvenile Myelomonocytic Leukemia: a Report from the Children’s Oncology Group. Cancer Discov 14, 1590–1598. 10.1158/2159-8290.CD-23-1376.

20. Wu, C., Xu, X., Shi, Y., Li, F., Zhang, X., Huang, Y., and Xia, D. (2024). Neutrophil Extracellular Trap Formation Model Induced by Monosodium Urate and Phorbol Myristate Acetate: Involvement in MAPK Signaling Pathways. Int J Mol Sci 26. 10.3390/ijms26010143.

